# An interlaboratory study of complex variant detection

**DOI:** 10.1101/218529

**Authors:** Stephen E Lincoln, Justin M Zook, Shimul Chowdhury, Shazia Mahamdallie, Andrew Fellowes, Eric W Klee, Rebecca Truty, Catherine Huang, Farol L Tomson, Megan H Cleveland, Peter M Vallone, Yan Ding, Sheila Seal, Wasanthi DeSilva, Russell K Garlick, Marc Salit, Nazneen Rahman, Stephen F Kingsmore, Swaroop Aradhya, Robert L Nussbaum, Matthew J Ferber, Brian H Shirts

## Abstract

Next-generation sequencing (NGS) is widely used and cost-effective. Depending on the specific methods, NGS can have limitations detecting certain technically challenging variant types even though they are both prevalent in patients and medically important. These types are underrepresented in validation studies, hindering the uniform assessment of test methodologies by laboratory directors and clinicians. Specimens containing such variants can be difficult to obtain; thus, we evaluated a novel solution to this problem in which a diverse set of technically challenging variants was synthesized and introduced into a known genomic background. This specimen was sequenced by 7 laboratories using 10 different NGS workflows. The specimen was compatible with all 10 workflows and presented biochemical and bioinformatic challenges similar to those of patient specimens. Only 10 of 22 challenging variants were correctly identified by all 10 workflows, and only 3 workflows detected all 22. Many, but not all, of the sensitivity limitations were bioinformatic in nature. We conclude that Synthetic controls can provide an efficient and informative mechanism to augment studies with technically challenging variants that are difficult to obtain otherwise. Data from such specimens can facilitate inter-laboratory methodologic comparisons and can help establish standards that improve communication between clinicians and laboratories.

## 1 INTRODUCTION

Next-generation sequencing (NGS) is a capable and cost-effective technique for detecting single-nucleotide variants (SNVs) and small insertions or deletions (indels) in relatively accessible parts of the genome [1]. Detecting such alterations is important in genetic diagnostics, and NGS has seen significant uptake in clinical laboratories for both germline and somatic DNA testing [2]. However, conventional NGS, like Sanger sequencing, has well-known limitations with other genetic variant types, including larger indels and complex alterations [3]. Furthermore, NGS can fail to detect variants in genomic regions that are not unique, are of low sequence complexity, or have very high or low GC/AT ratios [4,5]. Although technically challenging, variants of these types can be both medically important and prevalent among patients [6]. For example, in a companion study, we found that between 9% and 19% of pathogenic variants uncovered in patients were of types that are difficult for conventional NGS, depending on clinical indication [7].

Regardless, many published validation studies underrepresent or entirely omit these classes of variants. A recent systematic review found a “high degree of variability” among published validation studies, some of which did not even stratify performance by variant type (i.e. SNV versus indel) [8]. Indeed, the 2017 AMP (Association for Molecular Pathology) and CAP (College of American Pathologists) guidelines for NGS bioinformatics recommend that at least 59 variants of each type be included in validation studies [8]. One difficulty in accomplishing this is that few positive controls containing challenging variants in specific genes are readily available, despite the otherwise excellent results of the Genome in a Bottle (GIAB) and Genetic Testing Reference Materials (GeT-RM) programs [9–11].

In this interlaboratory study, we explored one potential solution to this problem: the creation of a synthetic reference sample (SRS) in which multiple technically challenging variants are synthesized and then introduced into a known human genomic background. This technique has previously been used in analyses of both germline and somatic panel tests [12–14], but these studies did not include many variants of high technical complexity. An approach using synthetic exogenous sequences has also been developed [15] but cannot be used with existing panel or exome tests. Moreover, all of these prior studies used a single NGS workflow (biochemistry and bioinformatics combined), although reference samples are maximally useful if they enable comparisons across a range of methodologies [16]. In this study, we created a single SRS with 22 challenging variants in commonly tested genes and evaluated it using 10 NGS workflows across an international group of 7 collaborating laboratories.

## 2 METHODS

We examined 80,000 clinical tests [7] performed by Invitae (San Francisco, CA), from which we selected 24 variants, all confirmed true positives and most both technically challenging and pathogenic, in 7 commonly tested tumor suppressor genes (Table 1). These variants were provided to SeraCare (Gaithersburg, MD), which synthesized plasmids containing these variants (Figure 1; Supplemental Methods). The plasmids were titrated into genomic DNA (gDNA) from the well-characterized GM24385 cell line [10] at concentrations to appear heterozygous.

DNA aliquots were provided to collaborating laboratories along with the list of genes of interest. The variant list was known only to SeraCare and non-laboratory staff at Invitae (SL, RT), who acted as data coordinators. Most laboratories used an Illumina (San Diego, CA) platform with hybridization-based targeting. One workflow (6) used whole-genome sequencing and another (8) used the Ion Torrent platform with AmpliSeq PCR-based targeting (ThermoFisher, Waltham, MA). Each workflow used a unique bioinformatics pipeline, including custom software, third-party software, and sequencing vendor–supplied software (Supplemental Methods).

## 3 RESULTS

Genetic variants in the SRS within the genes of interest included the following:

- 2 indels larger than 100 base pairs (bp)
- 6 mid-sized indels 11–28 bp
- 12 small indels less than 10 bp
- 4 deletions in short tandem repeats (STRs) of 3-4 nucleotide units with wild-type lengths 9–12 bp
- 1 larger tandem repeat expansion (2–3 x 24 bp)
- 2 homopolymer associated variants
- 5 deletion/insertion variants (delins)
- 3 variants in a segmental duplication
- 1 variant in an 80% GC region
- 2 SNVs near indels
- 1 SNV in which the genome and transcript references differed
- 15 additional benign SNVs in the 7 genes of interest

Note that some variants are counted in multiple categories in the list above. We considered 22 of these variants (Table 1) to present one or more technical challenges. Workflow 1A had initially uncovered these variants in patients and was primarily used to verify that they were present in the plasmid–gDNA mixture. Furthermore, we compared raw NGS data from this workflow with corresponding patient data, observing that the SRS presented technical challenges similar to those encountered in patient specimens. These challenges included biochemical artifacts, misalignments, clipped reads, and deviations from 50:50 allele fractions (Figure 1).

**Figure 1.**
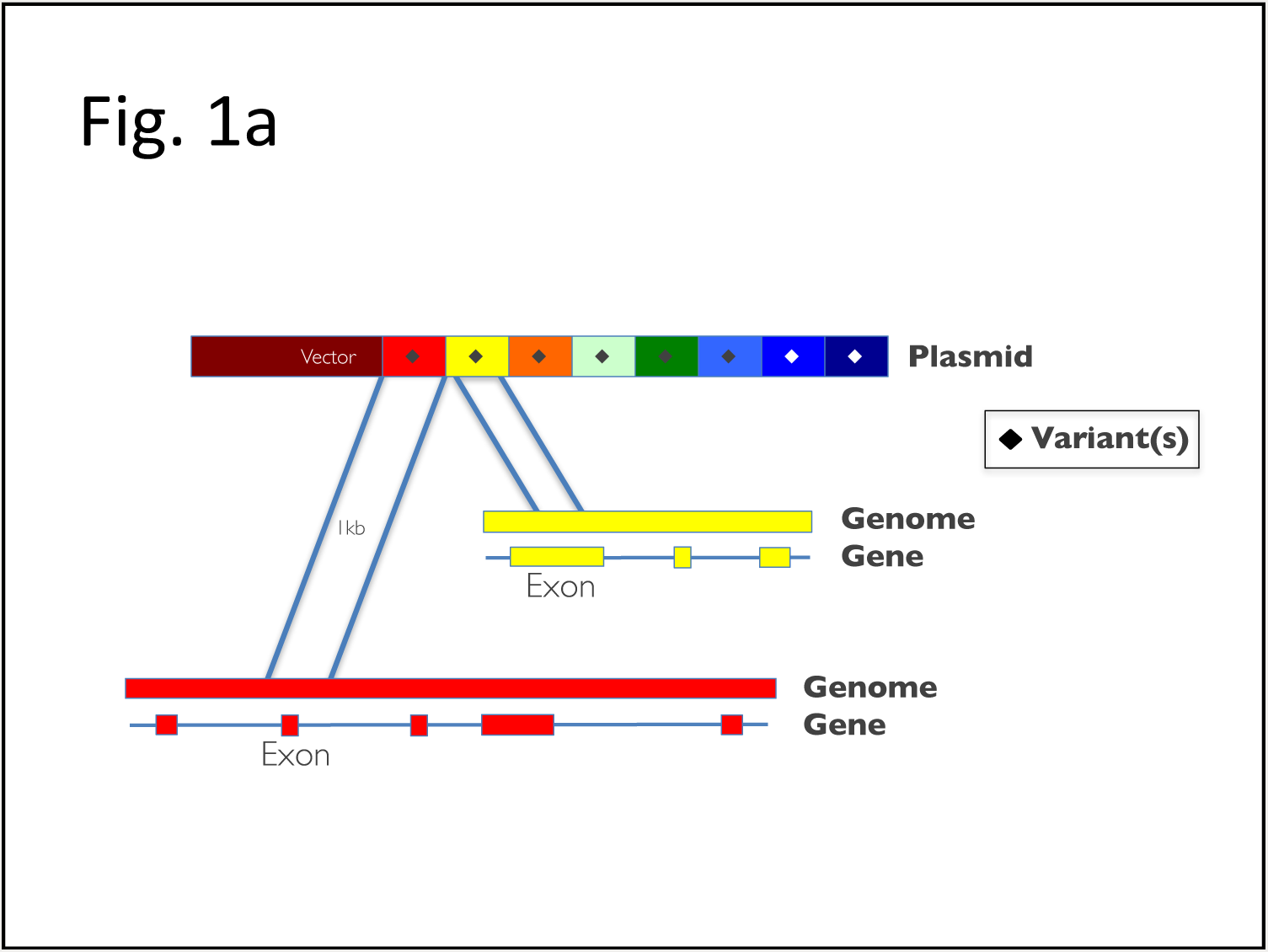
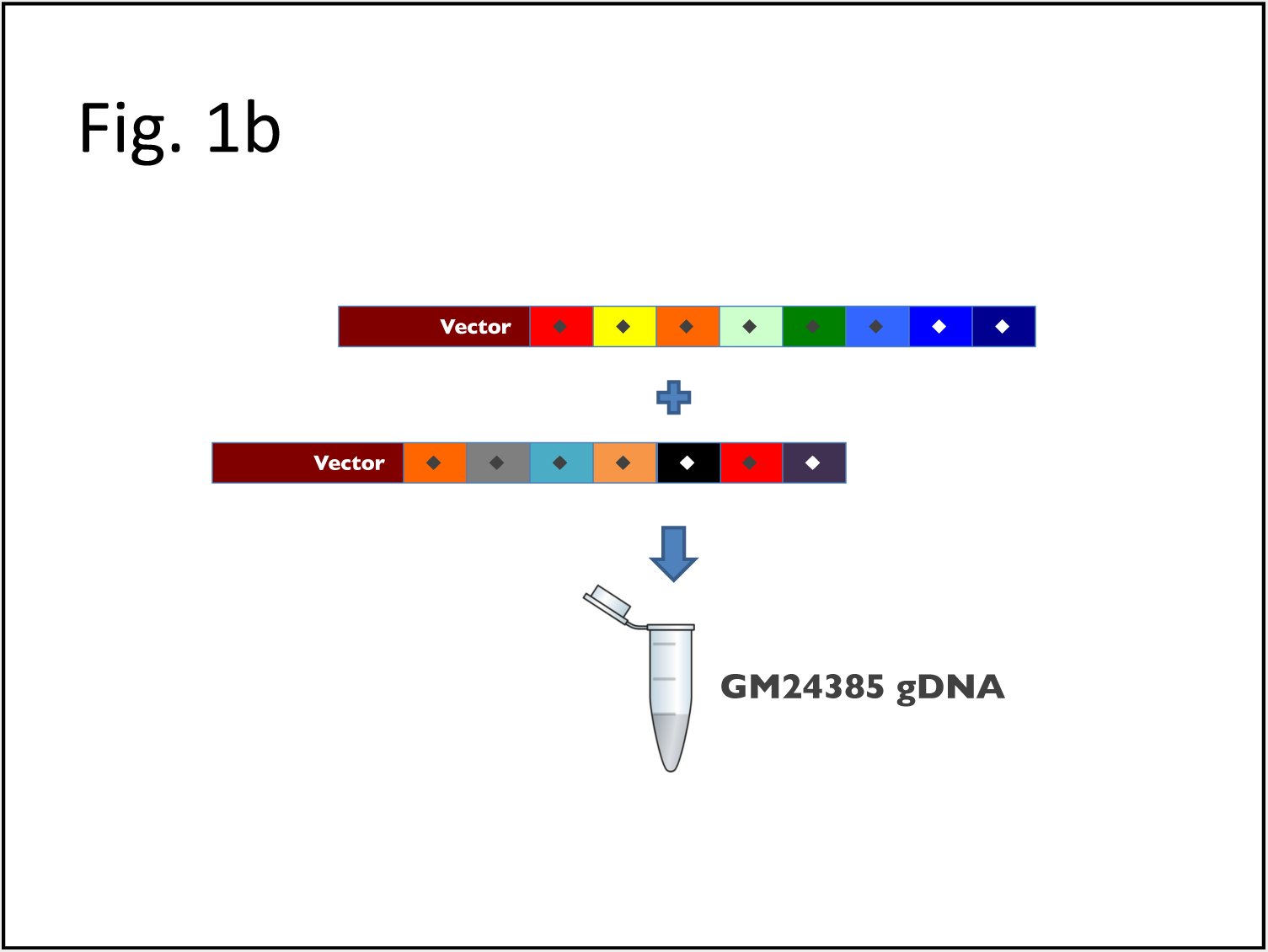
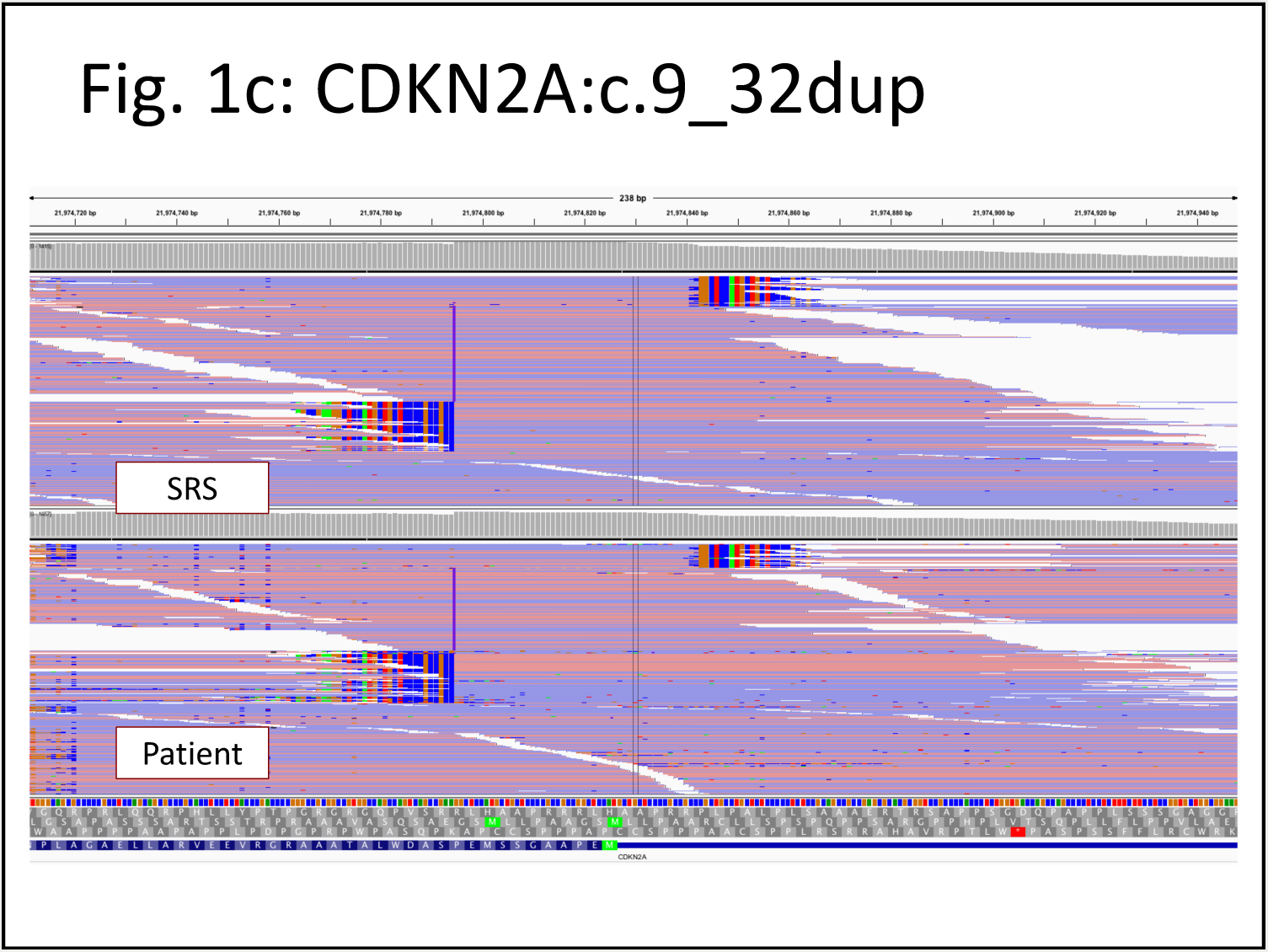
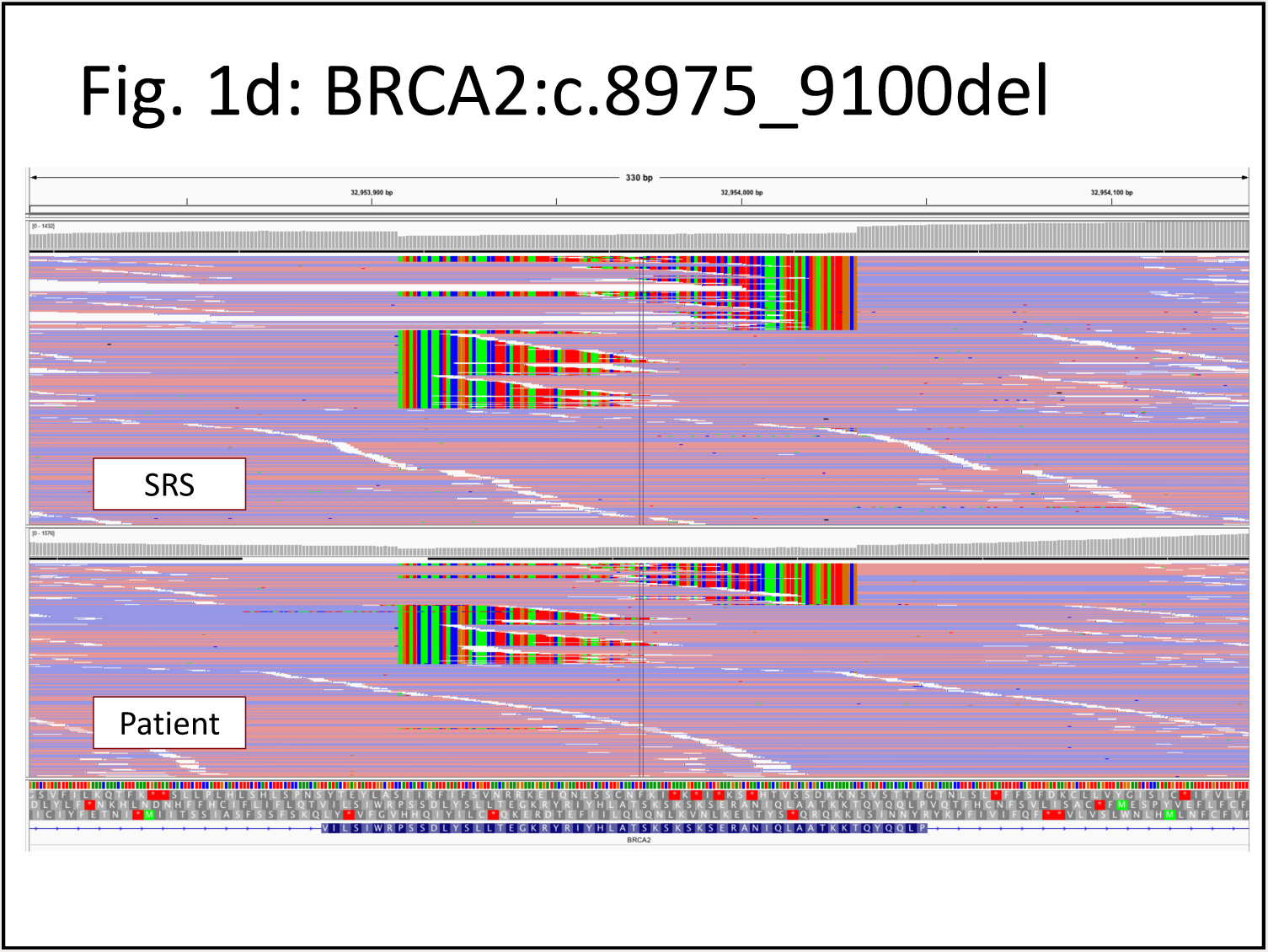
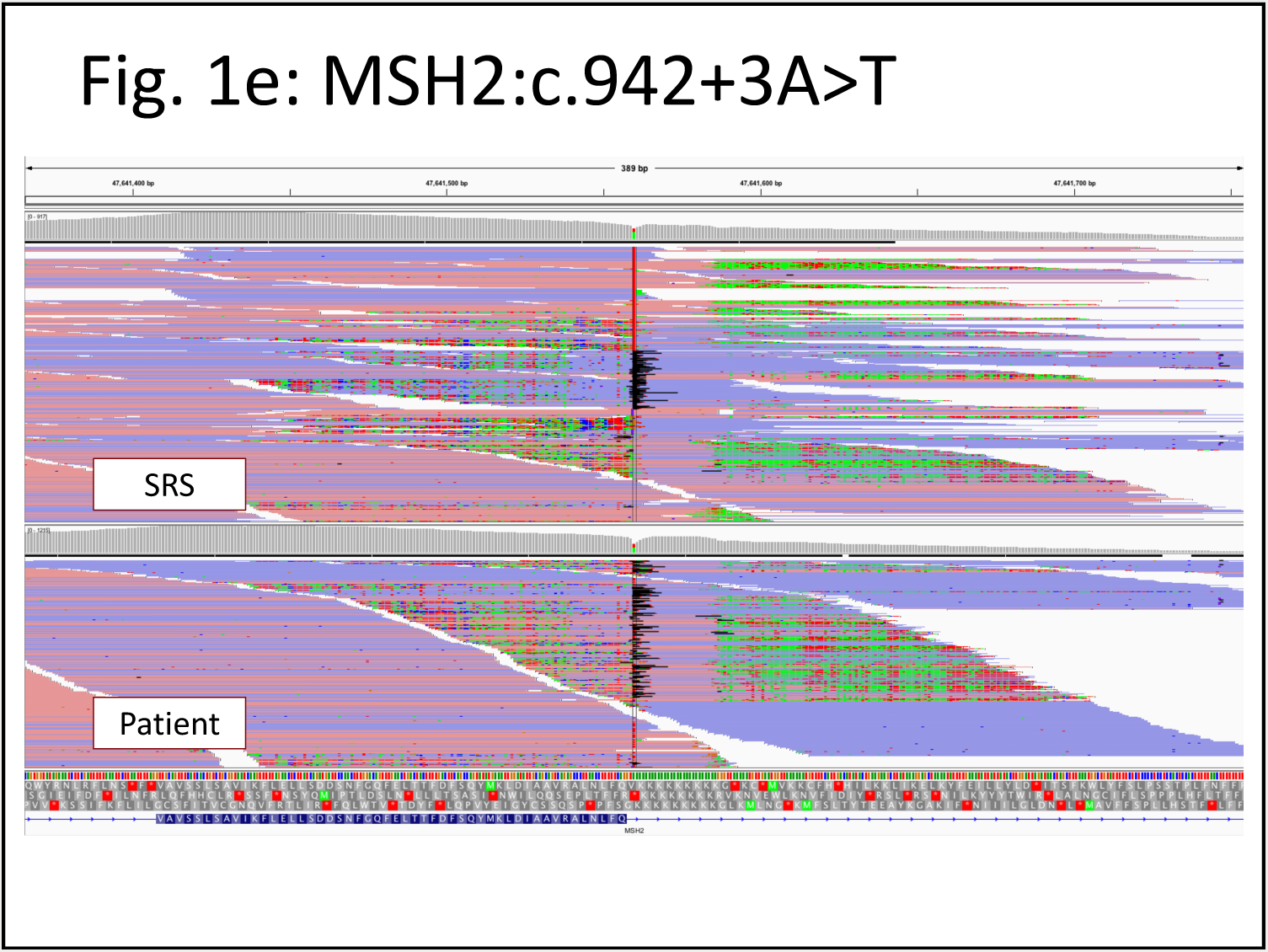
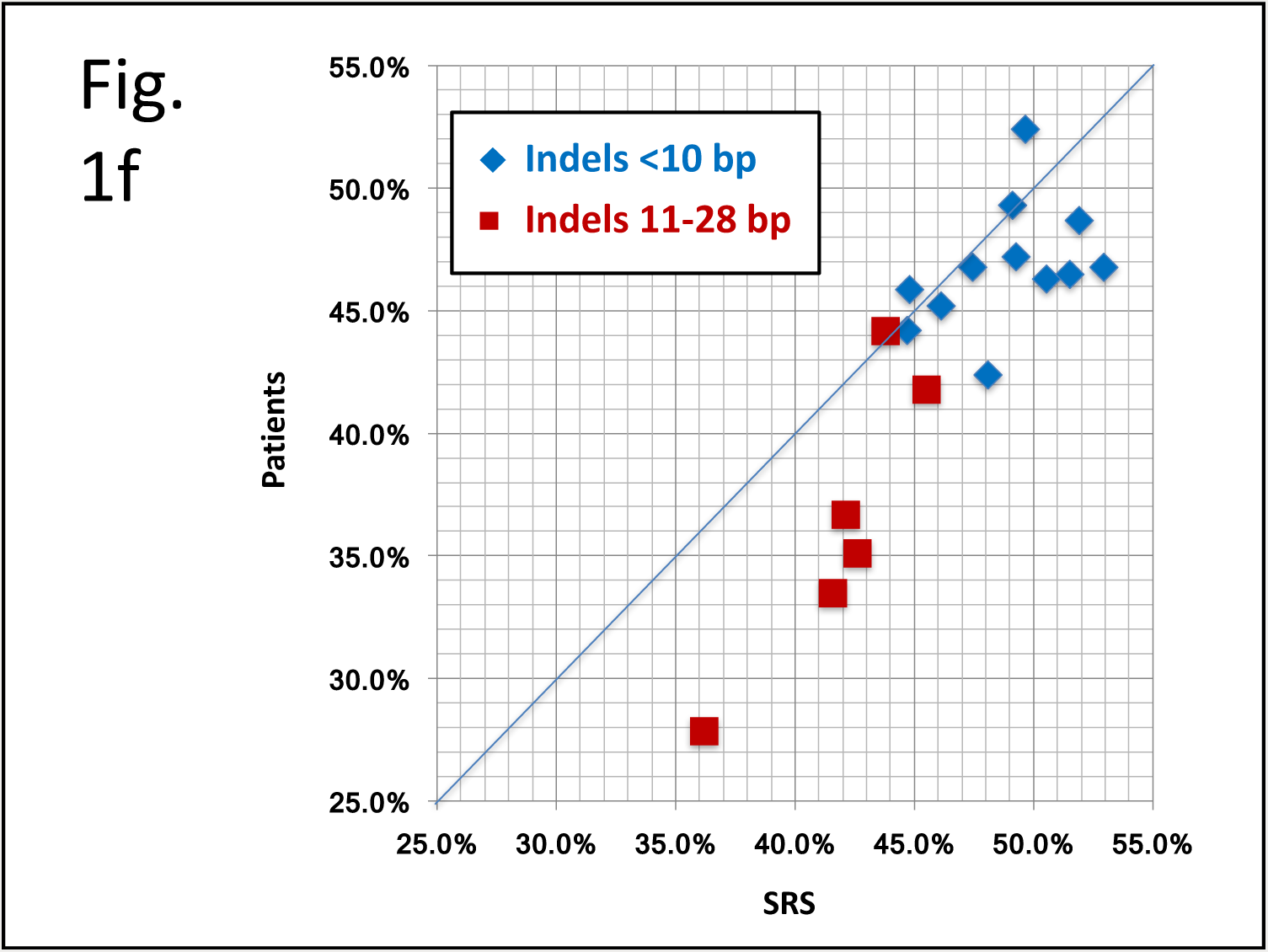
Construction and validation of the Synthetic Reference Sample (SRS). (a) Synthetic DNA fragments containing the desired variants with roughly 500 base pairs (bp) of flanking genomic sequence on each side were ligated into plasmid inserts. (b) Two such plasmids were titrated into human genomic DNA (gDNA) at concentrations appropriate for the plasmid variants to appear heterozygous against the gDNA background. When possible, we chose regions known to be homozygous in the background genome. (c–e) Next-generation sequencing (NGS) data for the plasmid–gDNA mixture was compared with that of representative patient specimens. Three variants are shown here (Table 1, variants N, L, and Q) each of which presents different technical challenges. (f) Variant allele fractions (VAFs) for small and midsized indel alleles observed in patient specimens were compared with VAFs observed in the SRS. Deviations from 50:50 are a known challenge with heterozygous indels owing to reference bias (i.e., NGS can be less efficient in both the capture and mapping of variant alleles compared with reference alleles), a phenomenon that becomes more pronounced with larger variants. The pattern of deviations is consistent between the SRS and patient specimens, although on average the SRS shows slightly higher VAFs (+3.2%). Compared with small variants, the midsized indels (11–28 bp) showed the lowest VAFs, as expected, and greater differences between the SRS and patients. The lowest VAF in patients (28%) and largest patient–SRS difference (+8.3%) were observed with variant R, a 28-bp deletion reaching from an exon into the neighboring intron. We did not include the 2 largest indels (>100 bp) in this graph, because the VAFs calculated by split-read analysis are not directly comparable to the VAFs of smaller variants. Qualitatively however, the VAFs for these 2 large events appeared even lower (<25%) in both the SRS and patients.

**Table 1.**
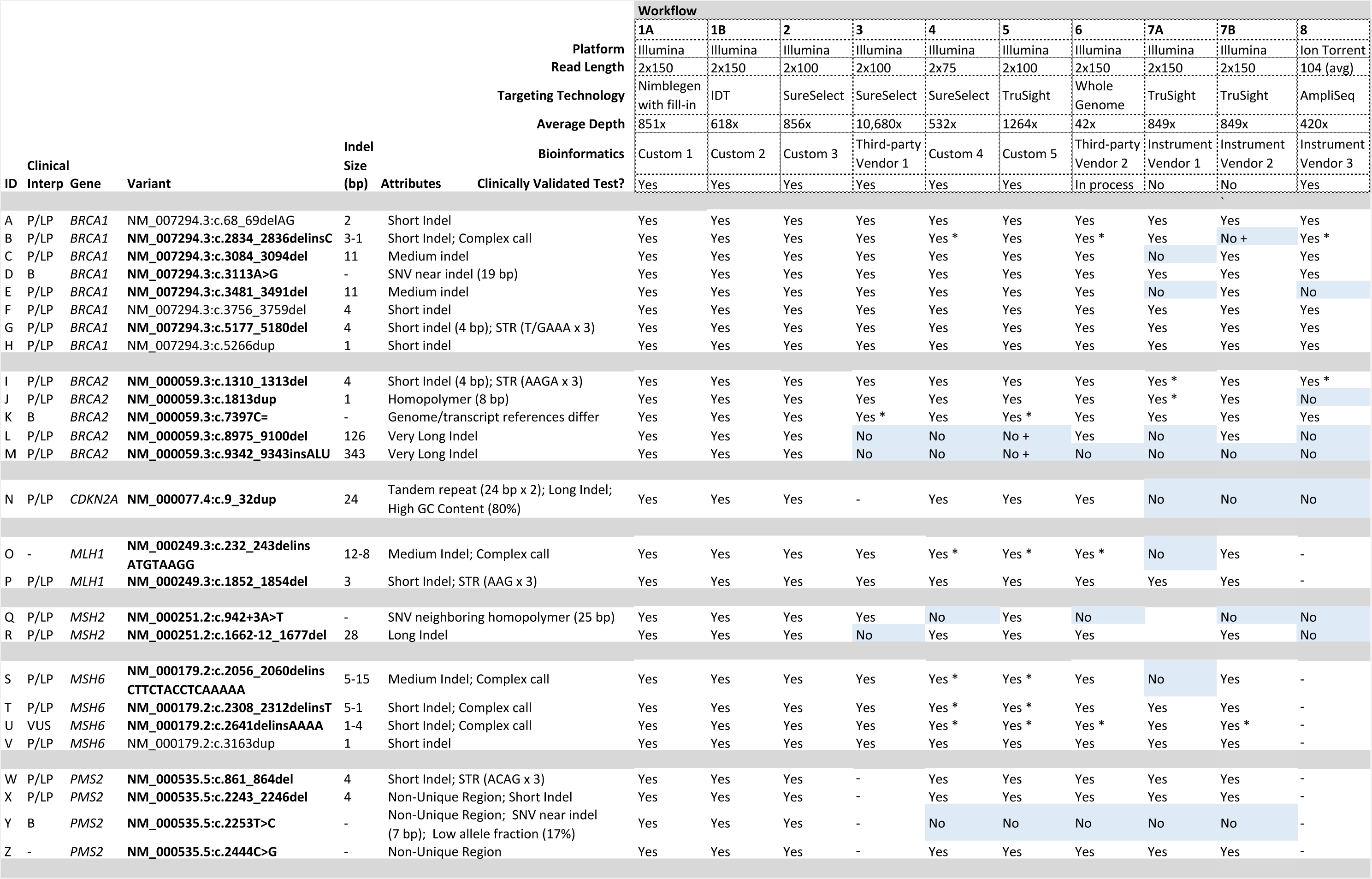
Results for each next-generation sequencing (NGS) workflow. “Yes” indicates that the variant was detected, “No” indicates that it was not. An asterisk (*) indicates that the variant was detected but described in a non-canonical form. Variants not detected include those with an incorrect call incompatible with the actual variant present (+), as well as those not reported in any form or filtered out (these have no other annotation). Blank cells indicate that the variant lies outside of the genomic regions interrogated by the workflow. The 22 variants in bold were considered technically challenging for reasons described in the attributes column. Clinical significance (interp) was extracted from ClinVar for all variants with a ClinVar entry. Abbreviations: B, benign; P/LP, pathogenic or likely pathogenic; VUS, variant of uncertain significance; bp, base pairs; Workflow 1A had previously detected all of these variants in patients and was primarily used to verify their presence in the synthetic control specimen. Workflow 1B is biochemically different than 1A and uses updated bioinformatics. Workflows 7A and 7B applied different bioinformatics pipelines to a single raw data set. All laboratories except number 3 reported that these sequencing depths were roughly typical for clinical testing, although they are high by research standards. Most laboratories did not confirm variants (e.g., with Sanger sequencing) in this study, as our focus was on NGS. Confirmation assays would likely have correctly resolved all of the incompatible variants shown here. A version of this table with additional information is provided as Supplemental Data. Additional information on the bioinformatics pipelines is available in the Supplemental Methods.

All 10 workflows detected all of the relatively “easy” SNVs and small indels within their target regions with one exception (Supplemental Data). However, only 10 of the 22 challenging variants were detected by all workflows, and just 3 workflows (including 1A) detected all 22. Nevertheless, most laboratories reported that some evidence of each variant was present in their raw NGS data, demonstrating that the SRS was compatible with the various biochemical methodologies used and suggesting that the sensitivity limitations observed were largely bioinformatic in nature.

Excluding workflows 7A and 8, discussed below, 3 bioinformatics errors were observed for the 18 small to mid-sized indels, including 1 incompatible call (delins variant B, worfkflow 7B in Table 1) and 2 false negatives: One of these (variant R, workflow 3) was related to the fact this 28 bp deletion starts 16 bp inside an exon and spans 12 bp into the neighboring intron. The other (variant N, workflow 7B) was in a complex tandem duplication in a low-complexity 80% GC region (this site had adequate coverage in this assay, however). All 4 small STR deletions were identified by all workflows. The large 126-bp deletion (L) was detected by all pipelines employing split-read algorithms [6,17,18] although 2 of those pipelines (6 and 7B) missed a large Alu insertion (M) also typically detected using split-reads.

Of the Illumina-based workflows, 7A and 7B showed the greatest limitations. These workflows used two different bioinformatics pipelines provided by the sequencing instrument vendor that were run with default parameters. Pipeline 7A omitted all indels larger than 10 bp, a limitation confirmed by the vendor’s support staff although we found no written documentation of it. Workflow 7B performed better than 7A although less well than workflow 6, which used the same targeting biochemistry (and shorter reads) but with custom bioinformatics.

Although workflow 8 (Ion Torrent AmpliSeq) was compatible with the SRS, the specific primers used for PCR-based targeting could not amplify 3 indel alleles (L, M, R) or left inadequate flanking sequences to align for 2 others (E, N) resulting in 5 false negatives due to allele dropout (Supplemental Figure 1). These issues would affect the same alleles in patients. This clinically validated workflow [19] is optimized for formalin-fixed specimens and thus uses small amplicons (mean, 104 bp), with each target site amplified by only 1 PCR primer pair, constraints that together resulted in these false negatives.

Workflow 8 also missed the 2 homopolymer-associated variants (J, Q), a known limitation of the Ion Torrent platform [1]. One of these variants (Q), an *MSH2* SNV neighboring a long homopolymer [7,20], was also missed by several Illumina workflows because of the simultaneous bioinformatics and biochemical artifacts it presents (Figure 1e). This one variant is both pathogenic and prevalent, comprising approximately 10% of positive findings in MSH2 [7].

Variants in *PMS2* exons 12–15 are challenging because of a pseudogene (*PMS2CL*) that prevents NGS reads from mapping uniquely. Workflows 3 and 8 ignored this region, whereas others correctly detected variants X and Z with the caveat that their locations (*PMS2* versus *PMS2CL*) could not be resolved without a separate assay. A heterozygous SNV in the base genome (Y) in this region was missed by most workflows: It appears in less than 20% of reads (because of the presence of *PMS2CL* and the plasmid) and is also near indel X, both known challenges for certain methods.

## 4 DISCUSSION

In this interlaboratory study, we created a single synthetic reference sample (SRS) containing 22 technically challenging variants—difficult to obtain otherwise in positive controls—in 7 commonly tested genes. We found that this sample is compatible with a diverse set of NGS biochemical methods and produces data that mimic those of patient specimens. These characteristics are crucial for such a sample to serve its intended purposes in methods development, optimization, comparison, and validation. The number of synthetic variants and genes in one SRS can be further expanded, making such samples even more useful and allowing the calculation of sensitivity confidence intervals by variant type, which we did not do because of the limited numbers of each variant type in our study.

Although not a proficiency test or formal benchmark, analysis of this SRS using 10 NGS workflows highlighted significant differences in sensitivity to certain variant types. Bioinformatics differences resulted in variable indel calling performance, particularly (but not exclusively) with larger variants. These issues were greatest with the sequencing vendor–supplied bioinformatics pipelines. In addition, multiple workflows failed to detect a pathogenic homopolymer-associated SNV, which presents both biochemical and bioinformatic challenges for NGS and which is explicitly mentioned as a common mutation in the ACMG technical standards for Lynch syndrome testing [20]. Furthermore, our amplicon sequencing method could not generate sequencing templates for many of the indels, resulting in allele drop-outs. (Other amplicon-based chemistries could potentially perform better, but we did not test any others in this study). These observations together reinforce the importance of carefully selecting biochemical methods, bioinformatics algorithms, and parameters for any NGS application.

Our study examined only sensitivity, not specificity, and trade-offs often exist between the two. Achieving high sensitivity for “hard” variants can result in additional false positives, which themselves may or may not appear hard. Laboratories should study this issue carefully using large sets of challenging variants if considering reducing the use of conformation assays even for “easy” SNVs or indels. Our study also focused on high-depth panel tests, although our 42x whole-genome data fared relatively well, reinforcing the fact that sequencing depth alone is an inadequate quality indicator [8]. Our study did not consider issues that arise in somatic testing even though the genes we investigated are included on many somatic panels and the variants we studied could influence therapeutic selection for PARP inhibitors or immunotherapies. A similar SRS for somatic tests could be constructed by titrating the same plasmids to a low variant allele fraction (VAF) as has been done previously [12–14]. The combined effect of reference bias (Figure 1f) with a low starting VAF would likely uncover additional sensitivity limitations for the medium and large indels, an important issue to explore further.

One complication in this study was the diversity of variant descriptions produced by different workflows for the same allele. Every delins variant was also reported as 2, 3, or 4 distinct neighboring variants by some workflows, usually without cis/trans phase information. We considered such calls a match as long as they appeared to describe the correct genomic sequence. However, this diversity presents significant challenges for variant databases, interpretation, and communication. We note, however, that some laboratories change variant descriptions after confirmation (e.g., Sanger sequencing), which was generally not used in this study.

Our SRS construction methodology has limitations. The largest low-complexity sequences we synthesized were 12-bp STRs and a 25-bp homopolymer, which are much smaller than, for example, the STRs underlying Huntington’s disease or fragile X syndrome. Our approach is also poorly suited to the construction of copy number variants, particularly deletions, for which genome editing may be superior. Indeed, we considered creating this SRS using genome editing, although we wondered whether cell lines with multiple mutations in DNA repair and cell cycle genes could be maintained. The variants we included in *PMS2* exons 12–15 proved useful, although current methods for disambiguating the location of such variants rely on long-range PCR or long read sequencing, both of which fail with the short fragments we synthesized. Longer fragments are possible but were not used here.

The data analysis for this SRS presented unique challenges. The junctions between fragments appear in split-read and paired-end analyses to be structural variant breakpoints (Supplemental Figure 2) that greatly outnumber signals from the 2 large indels in *BRCA2*. Laboratories can filter out these breakpoints by coordinate (Supplemental Data), and some did, although doing so can require modifications to a validated pipeline. Unsurprisingly, regions spanned by the plasmid also showed excess coverage and appeared as copy number gains by read-depth analysis. Both signals clearly indicate which exons may contain a synthetic variant, making it impossible to fully blind this SRS. Indeed, the higher than normal coverage at synthetic sites can aid in variant calling, making targeted downsampling of reads important for any formal benchmark (Supplemental Methods).

Despite these challenges and limitations, the collaborating laboratories in this study found that sequencing this SRS was highly informative and efficient. This single sample provided data on a wide range of variant types, and laboratories reported that this study uncovered new areas on which to focus future development. It is clearly impossible to include in validation studies every variant a sequencing-based test may encounter; however, it is crucial that an adequately large and representative set of variants be validated to exercise all aspects of the biochemical and bioinformatics procedures of a test [8]. Indeed, this single sample would greatly increase the number of non-SNV variants included in many validation studies [7]. Such specimens not only can help improve methods used by individual laboratories but also can improve transparency and communication about test limitations among laboratories and their end users, particularly non-geneticist clinicians.

## Acknowledgements

The authors thank Nancy Jacoby (Invitae) and Sarah Munro (National Institute of Standards and Technology) for help with the manuscript. We also thank Simon Cawley (Thermo Fisher) for discussion of the AmpliSeq results, and the Illumina staff for discussion of the MiSeq Reporter and BaseSpace results.

## Disclosures

Authors RG, CH, and FT work for SeraCare, the company that created the synthetic reference sample used in this study and provided it to the collaborating laboratories at no cost. SeraCare also provided sequencing reagents used in this study to the National Institute of Standards and Technology (NIST). No other financial relationship exists between the authors or their institutions and SeraCare. In addition, Thermo Fisher provided sequencing reagents for this study to NIST.

Authors SA, SC, WD, AF, MF, EK, SK, SL, SM, RN, NR, BS, SS, and RT work for laboratories that provide clinical genetic testing services. SA, SL, RN, RT, and RG own stock and/or stock options in their respective employers. SL owns stock in Illumina and Thermo Fisher. MF and EK receive royalties from SoftGenetics. NR serves as a consultant to AstraZeneca, and MF serves as a consultant to OneOme.

Certain commercial equipment, instruments, or materials are identified in this paper only to specify the experimental procedure adequately. Such identification is not intended to imply recommendation or endorsement by the National Institute of Standards and Technology, nor is it intended to imply that the materials or equipment identified are necessarily the best available for the purpose.

## Funding

Authors SA, MC, WD, AF, MF, CH, EK, RG, SL, RN, MS, FT, RT, PV, and JZ were funded by their respective employers. SK, SC, and YD were funded by NICHD/NHGRI grant U19HD077693. NR, SM, and SS were funded by Wellcome Trust grant 200990/Z/16/Z. BS was funded by the University of Washington and grants from the Damon Runyon Cancer Research Foundation (DRR-33-15) and the NIH (R21HG008513, NCI 5P30 CA015704-39).

